# Addressing technical pitfalls in pursuit of molecular factors that mediate immunoglobulin gene regulation

**DOI:** 10.1101/2024.03.06.582860

**Authors:** Eric Engelbrecht, Oscar L. Rodriguez, Corey T. Watson

## Abstract

The expressed antibody repertoire is a critical determinant of immune-related phenotypes. Antibody-encoding transcripts are distinct from other expressed genes because they are transcribed from somatically rearranged gene segments. Human antibodies are composed of two identical heavy and light chain polypeptides derived from genes in the immunoglobulin heavy chain (IGH) locus and one of two light chain loci. The combinatorial diversity that results from antibody gene rearrangement and the pairing of different heavy and light chains contributes to the immense diversity of the baseline antibody repertoire. During rearrangement, antibody gene selection is mediated by factors that influence chromatin architecture, promoter/enhancer activity, and V(D)J recombination. Interindividual variation in the composition of the antibody repertoire associates with germline variation in IGH, implicating polymorphism in antibody gene regulation. Determining how IGH variants directly mediate gene regulation will require integration of these variants with other functional genomic datasets. Here, we argue that standard approaches using short reads have limited utility for characterizing regulatory regions in IGH at haplotype-resolution. Using simulated and ChIP-seq reads, we define features of IGH that limit use of short reads and a single reference genome, namely 1) the highly duplicated nature of DNA sequence in IGH and 2) structural polymorphisms that are frequent in the population. We demonstrate that personalized diploid references enhance performance of short-read data for characterizing mappable portions of the locus, while also showing that long-read profiling tools will ultimately be needed to fully resolve functional impacts of IGH germline variation on expressed antibody repertoires.

## Introduction

The adaptive immune system employs B cell receptors (BCR) to mount effective responses to antigens. Variation in the BCR/antibody (Ab) repertoire has been linked to differential immune responses in a variety of disease contexts including infection, cancer, and autoimmunity [1–6]. This has prompted interest to better understand the development of the BCR repertoire and factors that contribute to variation in the circulating Ab repertoire in both healthy and disease states [1,7–12]. BCRs and Abs are composed of two identical heavy and light chains, which are encoded by genes at the immunoglobulin (IG) heavy (IGH) and light chain (kappa, IGK; lambda, IGL) loci. The genes within the IG loci serve as a primary source of the tremendous diversity observed among expressed Abs. Across the three primary IG loci, there are > 160 phylogenetically related functional/open reading frame variable (V), diversity (D) (in IGH), and joining (J) genes. In developing B cells, V, D, and J gene segments are recombined such that a single gene of each segment type is selected to form a rearranged gene that encodes the variable domain of the heavy (VDJ) or light (VJ) chain of an Ab. Diversity in antigen-naïve B cell Ab repertoires results from combinatorial and junctional diversity from V(D)J recombination, as well as the effective pairing of heavy and light chain segments [13].

IGH V(D)J gene rearrangement is mediated by the RAG complex (RAG1/2), which binds to recombination signal sequences (RSS) that flank IGHV, IGHD, and IGHJ genes to form the recombination center (RC). Most of our knowledge regarding specific factors involved in Ab repertoire regulation, including the process of V(D)J recombination, is derived from studies of inbred mice [14–24]. For example, the chromatin landscape of the C57BL/6 Igh locus is characterized by three loops anchored at boundaries of topologically associated domains (TADs) [20,25] associated with convergent CTCF binding motifs [23,26]. The probability of a gene being incorporated into the RC can be described as a function with multiple inputs, including 1) ability of RAG to bind the gene RSS 2) gene location within a TAD 3) spatial relationship of the gene to convergent CTCF motifs 4) gene-proximal DNA accessibility, and 5) binding of specific transcription factors that influence enhancer activity and enhancer-promoter contacts [14,15,19,20,25]. In addition, there are several well-described cis-elements in the C57BL/6 haplotype that regulate initial assembly of the RC: the intronic enhancer, Eμ, and the intergenic control region 1 (IGCR1) [22–24,26,27]. Critically, each of these factors that determine the Ab repertoire depend on interactions between germline DNA elements and DNA-binding proteins.

Studies in inbred mouse strains have provided a wealth of data on the molecular factors and mechanisms that orchestrate V(D)J recombination. These studies have examined V(D)J recombination during B cell development using short-read sequencing of chromatin domains and loops (*e.g.* HiC), accessibility (ATAC-seq, DNase-seq), as well as chromatin-associated architectural proteins (*e.g.* CTCF, Cohesin), histone modifications (*e.g.* H3K4me1, H3K27ac), and transcription factors (*e.g.* IRF4, PAX5) [15,19,20,26,28–30]. However, to date, these models have not systematically considered the impact of genetic variation that can occur between IG haplotypes [14]. In humans, haplotype diversity is a hallmark of the IG loci. Any two human haplotypes can differ by 100’s of kilobase pairs, driven by an enrichment of coding and non-coding single nucleotide variants (SNVs), short insertion-deletions, and structural variants (SVs) that can range from ∼9 - 270 Kbp and result in drastic changes in IG gene copy number [8,31–37]. In IGH, for example, 26 SV alleles have now been characterized, spanning >⅔ of IGHV genes [8]. If we are to fully understand the regulation of V(D)J recombination in humans, we need to develop strategies for assessing functional genomic data in outbred and genetically diverse populations.

In addition to inter-individual haplotype variation, the IG loci across species are enriched with segmental duplications (SDs) [33,38–40]. SD sequences are known blind-spots when mapping and analyzing short-read sequencing data [41,42] due to poor mappability. Indeed, we have demonstrated that short-read genotyping methodoligies for the IGH [8,36,43,44], IGL [37], and IGK [45] loci results in erroneous and missed variant calls. These pitfalls can be overcome through use of long-read sequencing and analysis frameworks that employ custom linear references [36,45].

There is mounting evidence that genetic variation in the IG loci is critical for establishing receptor diversity observed among human antibody repertoires [3,8,9]. This is highlighted by twin studies, which have shown that both naïve and antigen-stimulated Ab repertoires have heritable features [46]. In addition, IGH germline variants have been shown to directly impact Ab gene usage and antigen specificity [3,30–33]. Most recently, using matched AIRR-seq and haplotype-resolved IGH genotyping [36] in a cohort of 154 individuals, representing the largest study of this kind to date, we found that approximately half of common germline variants in IGH (minor allele frequency >5%) were associated with variation in the majority (73%) of IGHV, IGHD, and IGHJ gene usage frequencies in the IgM (naïve) repertoire [8]. Notably, intergenic variants were among those that exerted the largest effects on usage. Identification of these gene usage quantitative trait loci (guQTLs) provides a critical foundation for linking IGH germline variants to Ab repertoire diversity and will facilitate investigation of V(D)J regulation in human populations for the first time. Specifically, these data indicate that identified guQTLs likely illuminate cis-elements involved in DNA-protein interactions that regulate V(D)J recombination. A critical next step toward building the first models for understanding V(D)J recombination in human will require the integration of guQTLs with functional chromatin data (*e.g.*, ChIP-seq) to identify key factors underlying this complex molecular process and shed light on how diversity in the human repertoire is generated. This information will be critical for understanding differences in Ab-mediated responses in the human population in a variety of disease and clinical contexts.

Understanding how IG genetic variation impacts human Ab repertoires requires functional chromatin data accurately linked to human haplotypes. To date, these data have not been generated, and therefore mechanisms of V(D)J regulation in human populations are poorly defined. We propose that haplotype-resolved epigenomic data are critical for understanding V(D)J regulation in humans because the process of V(D)J recombination occurs on a single chromosome in developing B cells [47,48] and, as discussed above, current models of V(D)J recombination do not account for haplotype diversity. Here, we argue that the generation of haplotype-resolved regulatory maps for IGH cannot be accomplished using standard methods of chromatin profiling that involve mapping short reads to a single generic reference genome. We use simulated data derived from human IGH haplotypes to define locus and sequencing data characteristics that will hinder functional genomic studies in IG. These results clearly highlight the need for the development of novel informatics and molecular approaches for accurately identifying and characterizing molecular factors involved in V(D)J recombination at haplotype-resolution and at population-scale. Moving forward, we promote the adoption of new approaches that leverage the combined use of personalized reference sequences and long-read molecular assays.

## Results

### Segmental duplications hinder the accurate mapping of short reads in the IGH locus

Regulatory elements revealed by ChIP-seq peaks are primarily derived via mapping of short reads to a reference genome. Therefore, the identification of regulatory elements is dependent on accurate mapping of reads (*i.e.* to their original chromosomal positions). This bears consideration for portions of a genome enriched with repetitive or duplicated sequences that result in read mapping ambiguity [49]. Demonstrating this, random sampling of 500 genomic intervals (1.18 Mb in length) in the human reference GRCh38 resulted in a SD [50–52] content of ∼1% (**Figure 1A**), and it is estimated that ∼7% of sequence in a complete human genome is composed of SDs [42]. In contrast, ∼45% of the GRCh38 IGH locus (chr14:105860500-107043718) is composed of SD sequence (**Figure 1A**), highlighting the SD-rich nature of IGH that has been reported previously [32,33,53].

**Figure 1.**
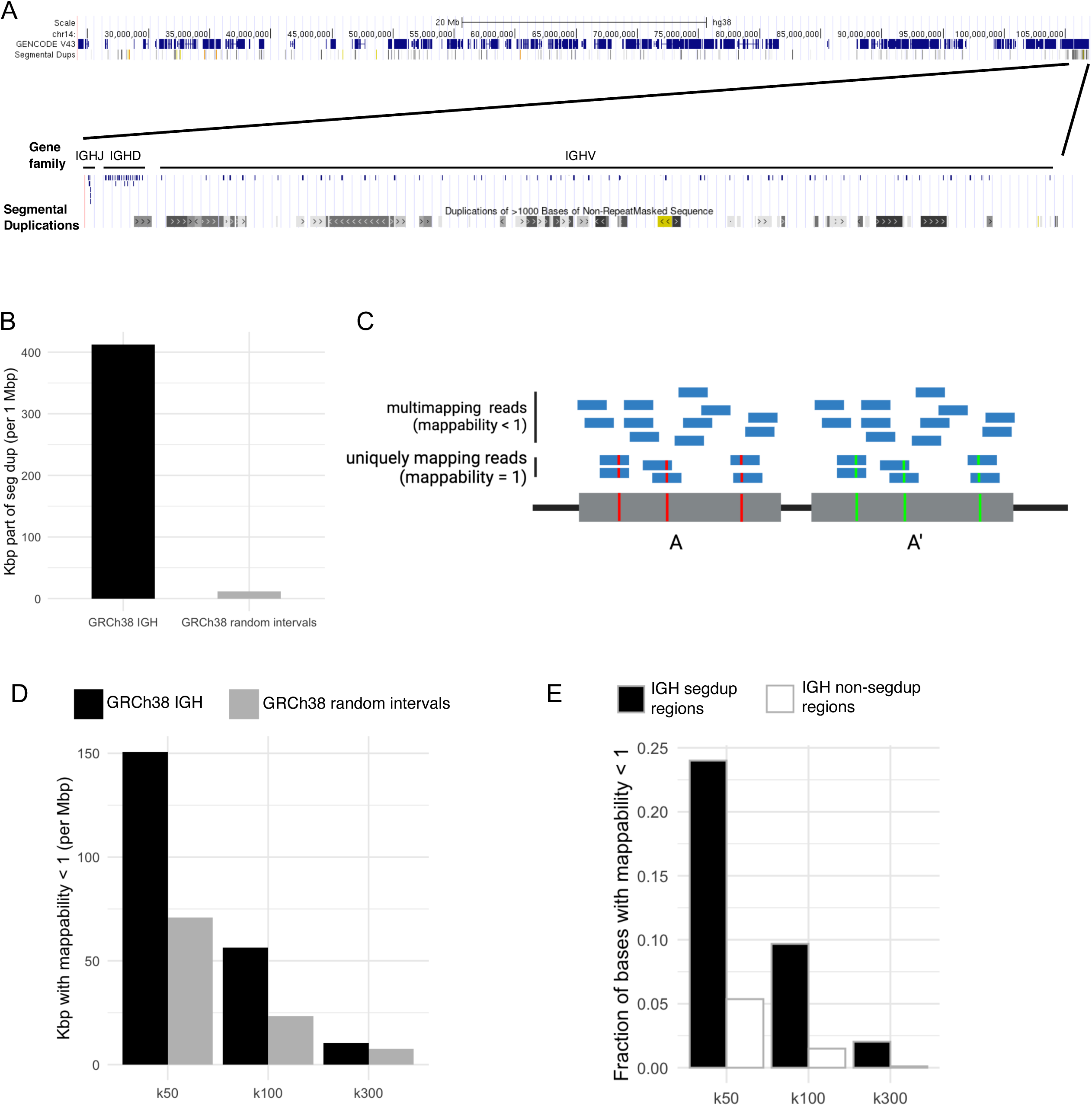
Segmental duplications in the IGH locus limit mappability. **(A)** UCSC genome browser (GRCh38) image depicting 85 Mb of chr14 including the IGH locus (top) and a zoomed-in view of the IGH locus with IGHJ, IGHD, and IGHV gene annotations (bottom). The segmental duplication track [50–52] is also shown. **(B)** Barplot of the number of Kbp within the IGH locus or random genomic intervals that overlap segmental duplication positions. **(C)** Scheme of a hypothetical segmental duplication with duplicates A and A’, which have single-nucleotide differences at highlighted positions. Multi-mapping (unmappable) reads align equally well to A and A’; mappable reads are more likely to overlap single-nucleotide differences. **(D)** Mappability of GRCh38 IGH depicting unmappable bases (per Mbp) for k-mers of 50, 100, and 300 bp. **(E)** Fraction of GRCh38 IGH that is unmappable for k-mers of 50, 100, and 300 bp. IGH positions are categorized according to overlap with segmental duplications.

SD regions in IGH present a problem when mapping short-read DNA sequencing data; a read derived from a SD region with 100% identity to the duplicate cannot be mapped to a unique position (**Figure 1C**). Given this, we would expect the accurate mapping of any given sequence k-mer to the IGH locus to be more challenging relative to the rest of the genome. To assess this assumption, we used mappability analysis to determine positions within IGH to which simulated reads (k-mers) could be uniquely mapped. Genomic positions with a mappability value of 1 are considered uniquely mappable, whereas regions with mappability values < 1 demarcate repetitive or duplicated regions [54]. For this analysis, we chose to use k-mer sizes of 50, 100, and 300 bp because 50 bp is the minimum recommended read length for multiple ChIP-seq assays (https://www.encodeproject.org/chip-seq/transcription_factor/), and 300 bp is the longest read length offered by commercially available Illumina platforms. We found that 50-mers, 100-mers, and 300-mers were not uniquely mappable in IGH for 13%, 5%, and 1% of the locus, respectively, corresponding to 150,710 bp, 56,266 bp, and 10,373 bp of unmappable sequence (**Figure 1D**). This is in contrast to the remainder of the GRCh38 assembly, for which the per-Mb mappability was approximately double that of IGH (**Figure 1D**). The fraction of SD sequences with mappability < 1 was more than 3-fold greater than the fraction of non-SD sequences with mappability < 1, highlighting the fact that SDs are enriched with unmappable (repeated) sequence (**Figure 1E**).

### Reads derived from large insertions absent from GRCh38 result in erroneous mapping

The results above demonstrate that even in a simple example for which reads are simulated directly from a single linear haploid reference, not all reads can be mapped with certainty to this same reference. This is critical, as it outlines a best-case scenario; however, this does not reflect mapping results that would be expected from short-read datasets derived from a diploid biological sample, particularly from an individual carrying SVs. We and others have demonstrated that the GRCh37 and GRCh38 human genome references do not effectively represent population-level IGH haplotype diversity, particularly with respect to SVs [8,33,36] (**Table 1**). For example, relative to the GRCh37 IGH assembly, many large insertions ranging from ∼9.5 kbp to ∼77.6 kbp have been identified in the human population (**Table 1**). These regions collectively harbor up to 19 IGHV genes, and represent ∼220 kbp and ∼164 kbp of sequence absent from IGH assemblies in GRCh37 and GRCh38, respectively. Importantly, many of these SVs harbor SDs in addition to those described in GRCh37 and GRCh38. This raises potential issues with the use of a single reference assembly for reliably mapping and interpreting functional genomic data from an individual carrying diverse IGH haplotypes. Specifically, DNA sequencing reads derived from insertions absent from a given reference assembly will either be mismapped to incorrect positions or unmapped, resulting in erroneous results or completely missing data. To assess the extent of this in IGH, we simulated 100 bp paired-end reads from 4 SVs (1 homozygous insertion; 1 heterozygous insertion; and 1 homozygous complex event) characterized from haplotype-resolved IGH assemblies for the 1KGP sample NA19240 (**Figure 2A**; [36], **see Methods**). Together, these SVs totaled ∼134.1 Kb of (haploid) sequence not represented in GRCh38 (**Figure 2A**). Reads derived from these insertions were mapped to GRCh38 using bowtie2 and parameters defined by ENCODE, which do not allow read multimapping (**Figure 2A, Figure S1**). We found that 100% and >98% of reads derived from the IGHV3-23D and IGHV3-30-region insertions mapped to the GRCh38 IGH locus, respectively (**Figure 2B-C**). Similarly, ∼86% of reads derived from the IGHV4-38-2 insertions mapped to GRCh38 IGH and ∼5% of reads were unmapped (**Figure 2B-C**). For the IGHV1-8 insertions, 30% of reads mapped to GRCh38 IGH. An additional ∼28% of reads from this insertion mapped to pericentromeric regions of chromosomes 15 and 16 (chr15:19999994-19987928, chr16:32,034,172-32,081,090, chr16:32,843,824-33,028,079, chr16:33,827,401-33,976,855) (**Figure 2B**), which harbor IGHV orphon genes [55–58]. In total, 77.9% of reads derived from insertions that were absent from GRCh38 were mapped to this genome reference. These data demonstrate that standard processing of short-read data from individuals with SV insertions via alignment to GRCh38 can result in false alignments of IGH sequences and thereby confound downstream interpretation.

**Figure 2.**
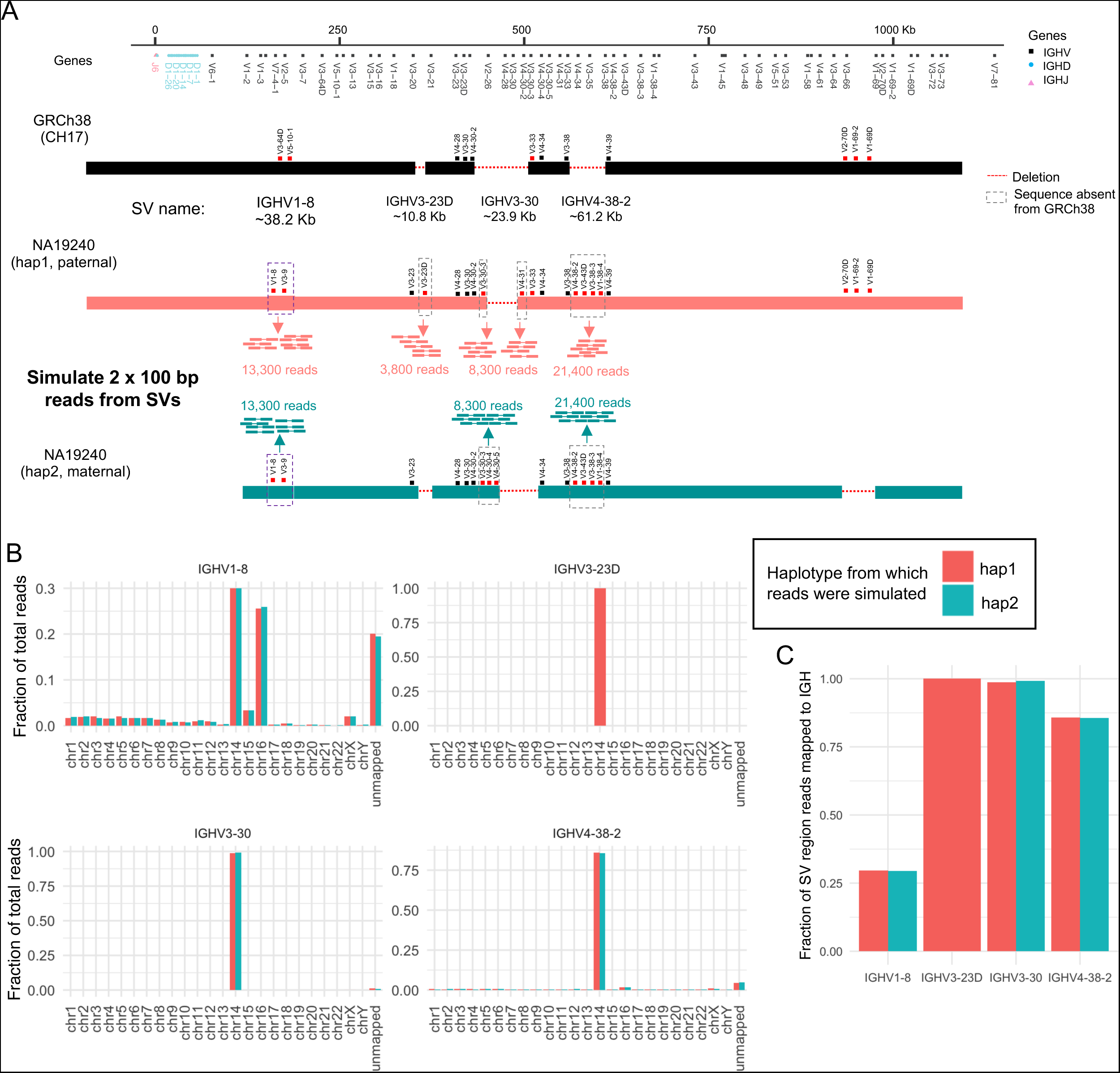
Structural variation limits use of a single linear IGH reference. **(A)** Schematic of a custom IGH reference (adopted from [8] and [36]) depicting inserted sequence absent from GRCh38 (below). Also shown are schematics of IGH haplotypes from the 1KGP sample NA19240. Red dashes indicate sequence that is absent relative to the custom IGH reference. Paired-end reads were simulated from two SV insertions for both haplotypes, a third (IGHV3-23D) SV insertion for only haplotype 1, as well as for a complex event, then reads were mapped to GRCh38 (see **Methods**). **(B)** Fraction of reads simulated off above-indicated insertions mapping to GRCh38 chromosomes, or unmapped. **(C)** Fraction of reads simulated off each SV region that mapped to GRCh38 IGH.

### Use of IGH-personalized diploid references allows for haplotype-specific mapping, but amplifies ambiguous mapping of short reads in homozygous and duplicated regions

The repetitive nature of IGH and presence of large SVs results in inaccurate short-read alignments. A possible solution could be utilization of a personalized genome reference [59]. This approach would provide the most relevant haplotypes on which to map an individual’s genomic data, and in theory allow reads to map specifically to their haplotype of origin. This ability to assess haplotype-specific signatures will ultimately be critical for assessing the role of genetic variants on V(D)J regulation. However, the use of diploid references, when both haplotypes are present, will also come with pitfalls. Specifically, an expansion of the overall mapping space will introduce increased chances for ambiguous mapping of short reads in homozygous regions, as well as in shared SDs within and between haplotypes (see **Figure S2**).

To assess the use of personalized diploid assemblies, we generated haplotype-resolved, IGH-personalized references for NA19240 and NA12878. Each personalized reference FASTA used GRCh38 as the backbone assembly, with an N-masked IGH locus (chr14:105860500-107043718) and two additional contigs representing the full-length assemblies of each IGH haplotype from either respective sample (**see Methods**). First, we determined the overall extent to which short reads can accurately map to an IGH-personalized reference, we performed mappability analysis on NA19240 and NA12878 IGH-personalized references using k-mers that correspond to read lengths commonly generated by short-read epigenomic assays (50 bp, 100 bp), and the upper-limit of commercially available Illumina sequencing (300 bp). Again, as with the assessment of GRCh38 as a reference, we found that read length was important for increasing overall mappability. For NA19240, mappability increased from ∼10% of the locus to ∼20% based on the use of 50-mers versus 100-mers, whereas the use of 300-mers resulted in ∼40% mappability (**Figure 3A**). A similar mappability pattern was observed for NA12878 (**Figure 3B**). Homozygosity impacted mappability to a significant extent as mappable regions were almost exclusively heterozygous when using 50-mers, 100-mers, and 300-mers (**Figure 3C**). Specifically, the number of homozygous bases in mappable regions ranged from 0 (NA12878 hap2 50-mer, 100-mer, 300-mer results) to 1860 bases (0.16% of IGH; NA19240 hap2, 300-mer result) (**Figure 3C**). Similarly, homozygous regions predominantly contributed to poor mappability (mappability < 1) regions when using 50-mers (85-90%), 100-mers (80-86%), and 300-mers (65-75%) (**Figure 3D**). Thus, these data demonstrate while, the use of personalized diploid assemblies, with both haplotypes present, can allow for haplotype-specific mapping in heterozygous regions, the use of short reads (100 bp), at best, can only accurately align to ∼20% of each IGH haplotype in an IGH-personalized reference. This indicates that longer reads will be required to fully capture haplotype-specific variation in the regulatory landscape.

**Figure 3.**
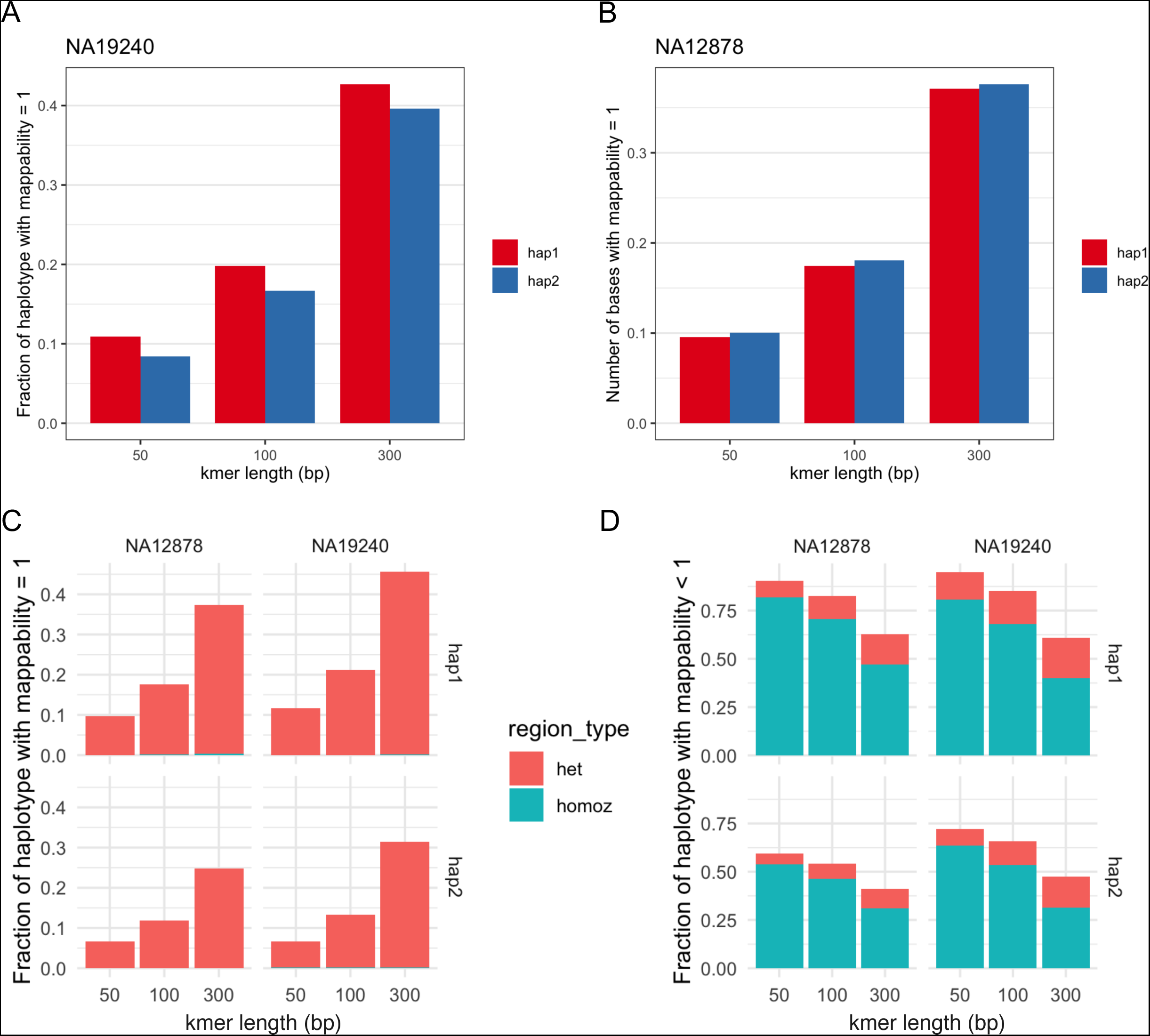
Limited mappability of IGH using short-read length k-mers for IGH-personalized genome references. **(A-B)** Mappability of IGH-personalized genome references for two samples (NA19240, NA12878) from the 1KGP. Fraction of IGH haplotypes with mappability=1 (uniquely mappable regions) were determined using simulated reads (k-mers) with lengths 50, 100, and 300 bp. **(C)** Fractions of indicated IGH haplotypes that are uniquely mappable with positions binned according to status as heterozygous or homozygous. **(D)** Fractions of indicated IGH haplotypes that are unmappable (mappability < 1) with positions binned according to status as heterozygous or homozygous.

To specifically address possible issues caused by assignments of short reads in SDs, we simulated 3,800 paired-end reads (100 bp) from the NA19240 hap1-specific duplication-insertion IGHV3-23D region (∼10.8 Kbp) and aligned these reads to the NA19240 IGH-personalized reference (**Figure S2**). We observed that 1,984 (∼52%) of reads mapped to their correct region-of-origin on hap1, while 878 reads (23.1%) mapped inaccurately to the IGHV3-23-region duplication block on the hap1 duplicated segment (IGHV3-23), and 938 reads (24.7%) mapped inaccurately to the IGHV3-23 region on hap2, which does not have the IGHV3-23D duplication-insertion (**Figure S2**).

### Alignment of short-read ChIP-seq to GRCh38 versus IGH-personalized references results in peak discrepancies

To directly assess the impacts of the mapping challenges outlined above on ChIP-seq peak calling, we aligned CTCF and H3K4me1 ChIP-seq reads from NA19240 (**Table 2**) to either an IGH-personalized reference or GRCh38. In the personalized reference, peaks in homozygous regions, which comprised 47% of hap1 and 40% of hap2 (**Figure 4A**), were assigned to both haplotypes. The peak set for hap1 consisted of 50 CTCF and 16 H3K4me1 unique haplotype-specific peaks. For hap2, we identified 70 CTCF and 24 H3K4me1 unique haplotype-specific peaks. This was in addition to 64 CTCF and 49 H3K4me1 peaks that were identified in homozygous regions and assigned to both haplotypes. In contrast, when we used GRCh38 as the reference, we identified 121 CTCF and 68 H3K4me1 peaks.

**Figure 4.**
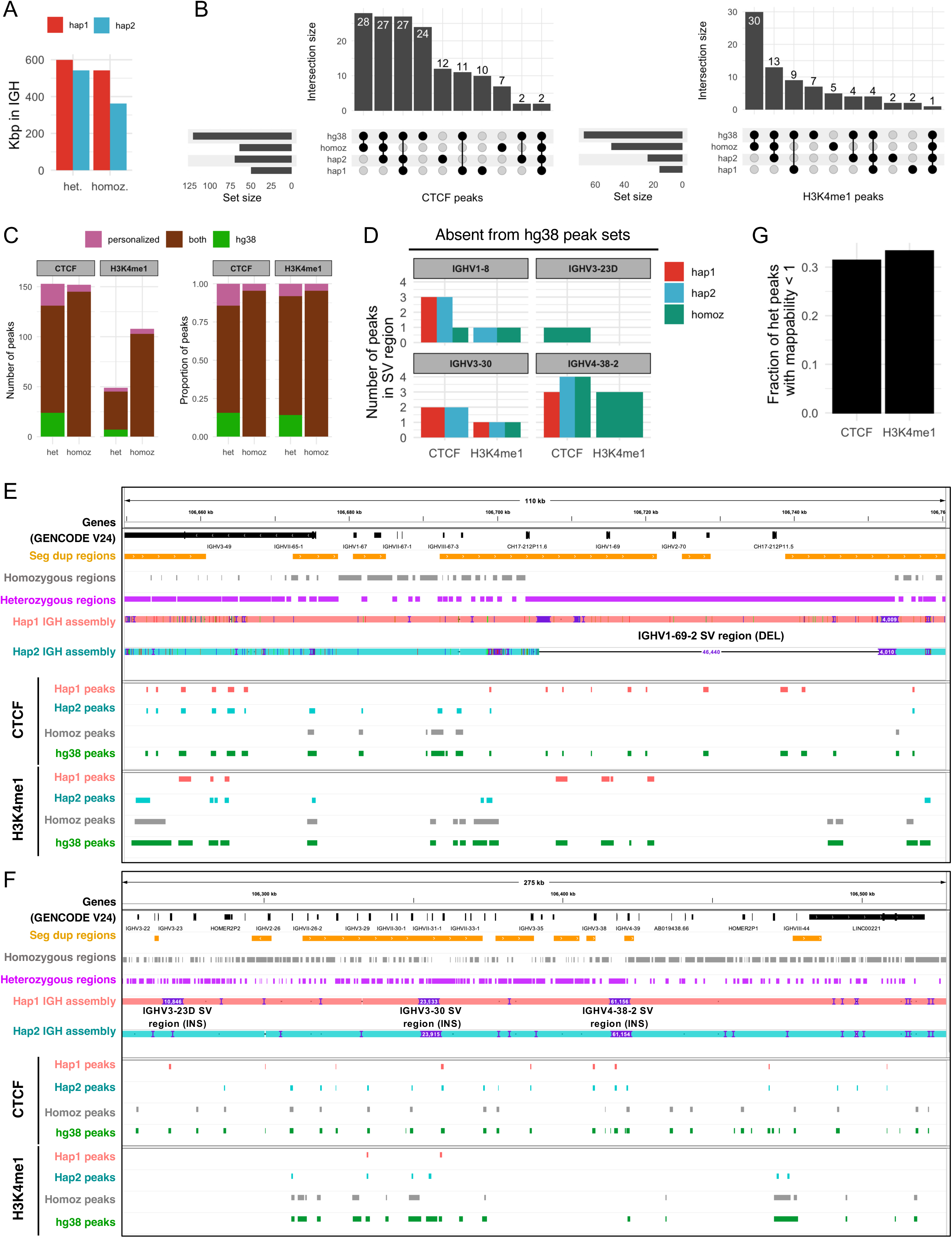
Alignment of short-read ChIP-seq data to an IGH-personalized genome reference versus GRCh38 results in discordant peak sets. **(A)** Number of bases that are within heterozygous or homozygous regions in the NA19240 IGH-personalized reference. **(B)** Upset plots depicting overlaps of ChIP-seq peaks for CTCF and H3K4me1 identified following alignment of reads to the IGH-personalized reference or hg38 (GRCh38). **(C)** Comparison of number of peaks (left) and proportion of peaks (right) resulting from alignment to the IGH-personalized reference or GRCh38 reference. “Personalized” indicates peaks uniquely assigned to hap1 and/or hap2 but not hg38. **(D)** Number of peaks within indicated SV regions for each haplotype; these peaks were not identified when reads were aligned to GRCh38. **(E-F)** IGV screenshots of regions that include the “IGHV1-69-2” SV region (**E**) and additional indicated SV regions (**F**). Also shown are gene annotations, segmental duplications [50–52], alignments of IGH hap1 and hap2 to chr14 of GRCh38, homozygous and heterozygous regions, and peaks for CTCF and H3K4me1 identified following alignment of reads to the IGH-personalized reference or hg38. **(G)** Fraction of peaks resulting from alignment to the IGH-personalized reference in heterozygous regions with low mappability (< 1).

To compare peak sets from the personalized reference to those called using GRCh38, we lifted over coordinates of peaks in each haplotype (hap1 and hap2) set individually to GRCh38. From these comparisons, we identified varying degrees of overlap between the peak call sets (**Figure 4B, 4C**). While the majority of peaks identified using GRCh38 were also found using the personalized reference, we noted that each peak set also contained unique peaks (**Figure 4B, 4C**). Specifically, alignment to GRCh38 resulted in 24 CTCF peaks and 7 H3K4me1 peaks (31 total) that were absent from the IGH-personalized reference peak set (**Figure 4B, 4C**). Conversely, 29 CTCF peaks and 9 H3K4me1 peaks (38 total) were identified uniquely in the IGH-personalized reference (**Figure 4B, 4C**). Among these 38 peaks, 31 (81.6 %) were identified within one of four SV regions discussed above (**Figure 4D, Figure S5**), which included 3 large insertions and a complex SV. By definition, these peaks could not be identified using GRCh38 because the SV sequences are not present in this assembly. Furthermore, it is likely that peaks unique to GRCh38 resulted from erroneous alignments of reads derived from these same four SV regions, as described above (**Figure 2**). We noted that the longest SV region containing IGHV4-38-2 (**Figure S3**) harbored more peaks than the other 3 SV regions (**Figure 4D**). We previously reported that the maternal haplotype (hap2) harbors a ∼46 Kbp deletion that includes *IGHV2-70D*, *IGHV1-69-2*, and *IGHV1-69D* [36]; accordingly, peaks in this region were assigned only to hap1 in the IGH-personalized analysis (**Figure 4E**). Additional examples of peaks assigned to hap1, hap2, and homozygous regions of the IGH-personalized reference and to the GRCh38 reference are shown in **Figure 4F**.

Having demonstrated that >50% of IGH haplotypes are not uniquely mappable using short read lengths (k-mers) (**Figure 3**), we sought to determine the number of peaks from the IGH-personalized reference peak sets that fell in unmappable regions. We considered only peaks in heterozygous regions, as peaks in homozygous regions cannot be assigned uniquely to either haplotype. Among 121 CTCF and 54 H3K4me1 heterozygous peaks, 38 (31.4%) and 18 (33.3%), respectively, fell in unmappable (mappability < 1) regions of the IGH-personalized reference (**Figure 4G**). Therefore, about ⅓ of peaks in heterozygous regions are within duplicated sequences (*e.g.* SDs) and cannot be confidently interpreted when mapping short-read ChIP-seq data to an IGH-personalized reference.

Taken together with our observations of erroneous mapping of reads derived from SV regions to GRCh38 (**Figure 2**), this analysis suggests that use of an IGH-personalized reference reduces erroneous mappings and offers potential for reads to map to their correct position. However, the potential for erroneous peaks persists in regions with poor mappability of short reads due to the presence of duplicated sequences in both homozygous and heterozygous regions.

### Long reads improve haplotype-resolved mappability for IGH-personalized genome references

Recent work has shown that long-read sequencing can haplotype-resolve epigenetic variation (*e.g.* CpG methylation, protein-DNA interactions) in complex genomic regions [60,61]. To assess the potential for long-read epigenetic assays to characterize IGH in a haplotype-resolved manner through use of an IGH-personalized reference, we determined the mappability of k-mers ranging from 1,000 - 25,000 bp for NA12878 and NA19240 (**Figure 5**). For NA19240, 5,000-mers were required to achieve 97% mappability for each IGH haplotype, and 12,000-mers increased mappability to 99% of the locus (**Figure 5A**). Relative to NA19240, IGH haplotypes of NA12878 were less mappable for each k-mer value, with only 87% and 93% of the locus being mappable using 5,000-mers and 12,000-mers, respectively (**Figure 5B**). This was likely explained by four long (>15 kb) stretches of homozygosity present in NA12878. This is in contrast to the NA19240 reference, which does not have any stretches of homozygosity >15 kb (**Figure S4**). Collectively, these data demonstrate that haplotype-resolved read mapping to an IGH-personalized reference with both haplotypes present will require departure from standard short-read epigenetic assays and implementation of novel, long-read based methods with read lengths of at least ∼8-16 kb in length.

**Figure 5.**
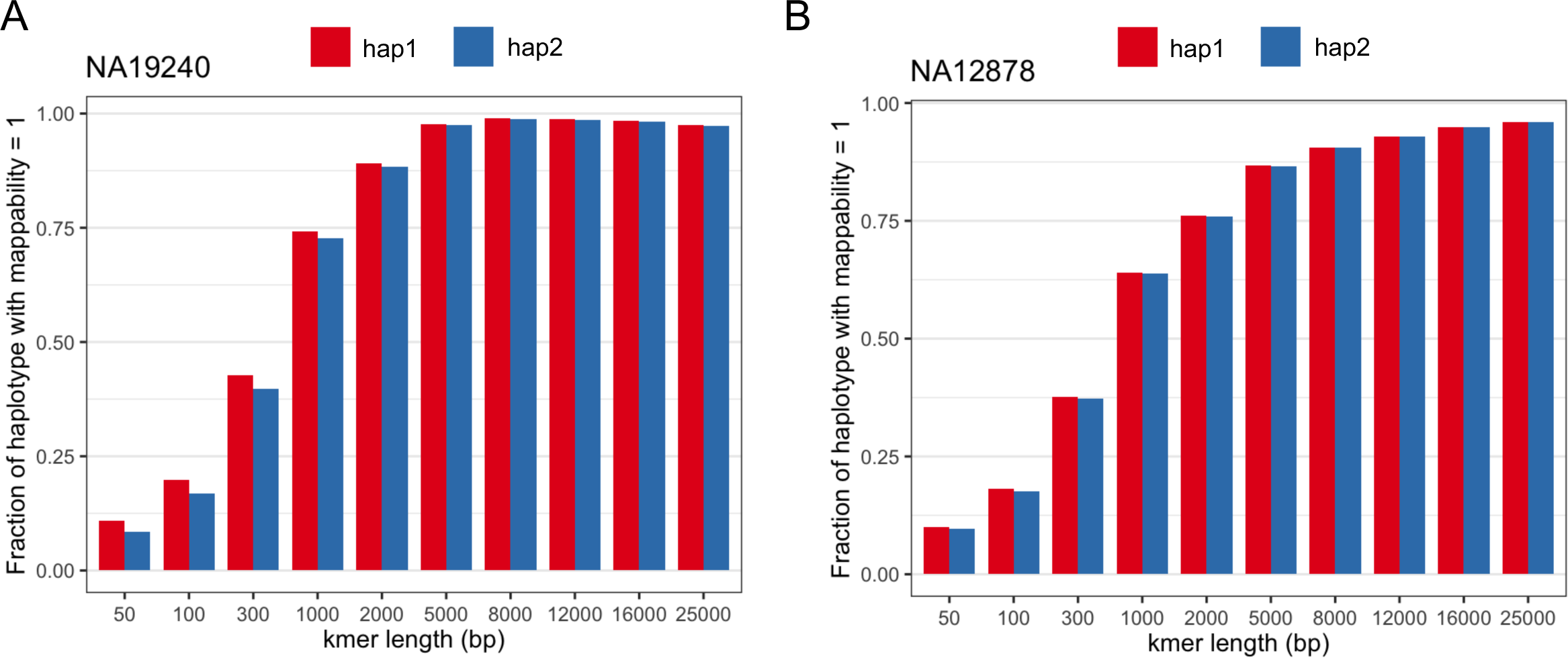
Long reads improve haplotype-resolved mapping to IGH-personalized genome references. **(A-B)** Mappability of IGH-personalized genome references for NA19240 and NA12878. Fraction of IGH haplotypes with mappability=1 (uniquely mappable regions) were determined using simulated reads (k-mers) with lengths ranging from 50 to 25,000 bp.

## Discussion

Mounting data in both human and model organisms indicate that V(D)J regulation and formation of an Ab repertoire are fundamentally linked to germline sequences within the IG loci [15,19,30]. However, there is a growing appreciation for the the fact that, unlike inbred mouse strains that carry homologous chromosome pairs with identical sequence (*i.e.,* absence of heterozygosity), humans inherit two distinct, and often genetically divergent, parental IGH haplotypes, either of which can undergo V(D)J recombination in a developing pro-B cell. This highlights a need to seek greater precision when analyzing functional genomic data in the pursuit of investigating factors that mediate V(D)J recombination. Specifically, this will require more comprehensive analyses in human populations at haplotype-resolution. Here, we have shown that in order to do this properly, we must consider the read lengths utilized by applied methods, the reference assemblies used for mapping, and also account for locus complexity and structure (*e.g.*, SDs and repeats), as well as SVs that occur at the population-level.

First, we demonstrated that when using a single linear reference, the mappability of short reads (50-300 bp; consistent with standard molecular genomic protocols) varied considerably depending on the length simulated. Nonetheless, in all cases this analysis indicated that a significant fraction of bases within a given IGH haplotype were inaccessible. As noted in previous studies [36,44], we highlight here that this is a result of the duplicated and repetitive architecture of the IGH locus. This initial simulation represented a best case scenario, but we showed that additional problems were encountered when considering reads derived from SVs absent from the reference assembly used for mapping. Specifically, all alignments to GRCh38 for reads derived from these SV regions are erroneous. Therefore, it is possible that short-read ChIP-seq peaks unique to the GRCh38 reference analysis may represent false signals secondary to erroneous mappings. Use of an IGH-personalized reference allowed short reads to map to SV regions that were absent from GRCh38; however, this approach still resulted in ∼30% of heterozygous peaks falling in regions with low mappability (k-mer 300 bp) due to duplicated sequences in the IGH-personalized reference. These data demonstrated that not accounting for such structural variation would result in a significant number of false-negative and -positive mappings.

Recent long-read based sequencing and assembly has opened up new opportunities to fully characterize IG haplotypes. With such assemblies in hand, molecular genomic data such as ChIP-seq could be mapped to personalized haplotypes improving mapping precision and data interpretation; this has been shown using genome-wide datasets [62]. However, within the IGH locus, we showed that when using short reads, the issues encountered because of SDs within a haploid reference remained, and in some cases were worsened when two haplotypes harboring shared SDs were used. Additionally, our analysis showed that, due to homozygosity, for a fraction of bases within IGH, short reads could not be mapped to specific haplotypes of the personalized reference.

Together, these results highlight that mitigation strategies are required when using short reads for molecular genomic studies in the IGH locus. One strategy could involve the tailoring of analysis to include only information gleaned from mappable regions of a given reference assembly, whether a generic haploid assembly or personalized reference is used. For example, one could permit downstream analysis of ChIP-seq for peak identification only for reads that align without mismatches to regions with mappability=1. However, in the case of a generic haploid reference assembly, the inclusion of as many known SVs in that reference would be ideal to account for haplotype variability in the donor from which experimental reads are derived. Additionally, if using a diploid personalized reference, regions of homozygosity could be collapsed for peak calling, in contrast to regions of heterozygosity in which haplotype-specific mapping is possible. Critically, these points should not be overlooked in non-human animal models. The mouse IG loci, for example, are also enriched for large duplications and repeats [63,64], and thus issues with read mappability would be expected.

Finally, our analysis indicated that the future could see significant improvements in molecular genetic studies of the IGH locus as assays that leverage longer reads are developed. Specifically, we showed that when using a personalized reference, the majority of bases (>95%) could be resolved (mappability=1) using reads >5 Kb. Single-molecule real-time (SMRT) long-read sequencing platforms from Pacific Biosciences (PacBio) are capable of detecting modified bases on native DNA. Oxford Nanopore Technologies (ONT), which reports DNA bases from changes in current across a pore-forming channel, can also detect modified bases on native DNA. Thus, already these technologies are being leveraged to measure DNA methylation, chromatin accessibility, and protein-DNA interactions on individual chromatin fibers ranging in length from ∼5 Kbp to ∼100 Kbp [61,65–69]. For example, ONT sequencing of a “haploid” human cell line (CHM13) yielded CpG methylation information resolved across SDs and complex structural events [42]. In addition, the technique “DiMeLo-seq” employs long-read (PacBio or ONT) sequencing to reveal protein-DNA interactions at base pair-resolution [61]. This method uses a fusion reagent composed of protein A and a nonspecific DNA adenine methyltransferase to “tag” DNA where a primary antibody (e.g. targeting CTCF) is bound to chromatin. Based on our results using simulated reads (**Figure 5**), we hypothesize that protein-DNA interactions can be haplotype-resolved for IGH using a technique such as DiMeLo-seq and an analysis framework that uses an IGH-personalized reference. Integration of guQTLs with accurately mapped epigenetic information will provide mechanistic insight into gene usage differences that associate with human haplotypes.

## Materials and Methods

### Segmental duplication analysis

The GRCh38 IGH locus was defined as chr14:105860500-107043718 (from IGHJ6 to the telomeric end, excluding the telomere). SD coordinates for GRCh38 were downloaded from the UCSC table browser (table: “genomicSuperDups”) [50–52]. SD coordinates from alternate contigs or overlapping blacklisted regions (https://github.com/Boyle-Lab/Blacklist/blob/master/lists/hg38-blacklist.v2.bed.gz) outside of IGH were filtered out from downstream analysis. SDs overlapping IGH were extracted using the bedtools (v2.30.0) ‘intersect’ tool.

The SD content of random genomic intervals (excluding IGH) was determined by first generating 500 random intervals equivalent in length to the IGH locus (1,183,218 bp) using the bedtools ‘random’ tool with options ‘-l 1183218 -n 500’. Total SD bases per interval were determined using bedtools ‘intersect’ with the -wo option. The sum of bases within the random intervals overlapping SDs was divided by the sum of random interval lengths, then multiplied by 1 million. Similarly, the total SD-overlapping bases in IGH was divided by the length of IGH, then multiplied by 1 million.

### Haplotype-resolved IGH assemblies for NA12878 and NA19240

NA12878 (proband) PacBio WGS HiFi reads were downloaded from GIAB (https://ftp-trace.ncbi.nlm.nih.gov/ReferenceSamples/giab/refsam_s3_urls) and NA19240 (proband) PacBio WGS HiFi reads were downloaded from HGSVC2 (http://ftp.1000genomes.ebi.ac.uk/vol1/ftp/data_collections/HGSVC2/working/20191005_YRI_PacBio_NA1924 <u>0_HiFi/</u>). Illumina WGS (30X coverage) for NA12891 and NA12892 (parents of NA12878) and NA19239 and NA19238 (parents of NA19240) were downloaded from the IGSR (https://www.internationalgenome.org/data-portal/sample). Additional HiFi (Sequel IIe system, Pacific Biosciences) for each trio were generated using IGH-capture and IGenotyper as described previously [36].

HiFi reads from each proband were aligned to a previously described reference [8,36] that is publicly available (https://github.com/oscarlr/IGenotyper#getting-igh-specific-reference) using minimap2 (v2.26-r1175) with the option ‘-ax map-hifi’. Notably, this reference includes inserted sequences for the SV regions termed “IGHV1-8”, “IGHV3-23D”, “IGHV3-30”, and “IGHV4-38-2”. Non-supplementary and non-secondary alignments to IGH were extracted using samtools (v1.9) ‘view’. Haplotype-resolved IGH assemblies were generated using hifiasm (v0.18.2-r467) with the trio option. Small gaps in the hifiasm-generated assemblies were filled in using published phased IGH contigs generated by IGH-capture and IGenotyper for NA12878 and NA19240 [36] with a custom python script. For each proband, an IGH-personalized reference was generated by N-masking the IGH locus of GRCh38 (chr14:105860500-107043718) and appending each IGH haplotype as an entry in the FASTA file.

To validate the IGH-personalized references, parental IGH-capture PacBio reads were aligned to IGH-personalized references using minimap2 with the option ‘-ax map-hifi’ and phased using whatshap (v1.7.dev54+g0cb3793) [70]. For each proband, IGH-capture and WGS HiFi reads were aligned to their respective IGH-personalized reference. The resulting alignments of trio data to each IGH-personalized reference were manually checked in the Integrative Genomics Viewer (IGV) [71] for assembly (e.g. phase-switch) errors. Haplotype-resolved NA12878 and NA19240 IGH assemblies are available at <u>10.5281/zenodo.10674652</u>.

To determine heterozygous positions in diploid IGH assemblies, IGH haplotypes were aligned using minimap2 with the option ‘-ax asm20’ and variant positions (SNVs, insertions and deletions) for each haplotype were determined using a custom python script (<u>10.5281/zenodo.10674652</u>). These variant positions were extended bidirectionally by 49 bp (50-mer analysis), 99 bp (100-mer analysis), or 299 bp (300-mer analysis) bp using bedtools ‘slop’ and the resulting heterozygous intervals were merged for each using bedtools ‘merge’. To generate the NA19240 IGH-personalized reference for ChIP-seq analysis (**Figure 4**), the 300-mer parameter was used. Homozygous regions, defined as all non-heterozygous intervals for each IGH haplotype, were generated using bedtools ‘subtract’.

### Alignment of NA19240 SV sequences to GRCh38

IGH gene coordinates were lifted over from a custom IGH reference [8,36] to each IGH haplotype of NA19240 using custom python scripts. The GRCh38 IGH haplotype (chr14:105860500-107043718) was aligned to each IGH haplotype of NA19240 separately using minimap2 with options ‘-x asm20 -L -a’. Coordinates of breaks in the alignment of the GRCh38 IGH haplotype to each NA19240 IGH haplotype are shown in **Figure S1**; these coordinates were input to the tool bedtools ‘getfasta’ to extract sequences. Paired-end reads were simulated from these sequences using the readSimulator.py python script (https://github.com/wanyuac/readSimulator) with parameters ‘--simulator wgsim --iterations 1 --readlen 100 --opts ’-e 0 -r 0 -R 0 -X 0 -h -S 5’. The simulated reads were aligned to GRCh38 using bowtie2 with the ‘-X2000’ option as described for processing of paired-end ChIP-seq data by ENCODE (https://github.com/ENCODE-DCC/chip-seq-pipeline2/blob/master/src/encode_task_bowtie2.py). Read mapping statistics were determined by parsing BAM files using Rsamtools (https://github.com/Bioconductor/Rsamtools).

### Mappability analysis

Genmap (v1.3.0) [54] ‘index’ was run on GRCh38 and IGH-personalized references for NA19240 and NA12878. Mappability of these references was determined using the Genmap ‘map’ tool with options ‘-T 10 -E 0 -K <INT> -bg’, with <INT> values of 50, 100, 300, 1000, 2000, 5000, 8000, 12000, 16000, and 25000.

### ChIP-seq data

Replicates of CTCF and H3K4me1 ChIP-seq from the NA19240 LCL were downloaded from SRA accessions listed in **Table 2**.

### ChIP-seq analysis

Homozygous regions of the NA19240 300-mer IGH-personalized reference (described above) were N-masked on hap2 to permit mapping to only hap1. Reads were aligned to the NA19240 IGH-personalized reference or GRCh38 using bowtie2 with parameters --very-sensitive --non-deterministic. Replicate BAM files were input for peak calling with Genrich (https://github.com/jsh58/Genrich) using parameters “-s 0 -t <replicates> -r -v -y -q 0.05”. Peaks in homozygous regions were identified using bedtools ‘intersect’ with parameters “-wa -a [<Hap1_peaks.bed> or <Hap2_peaks.bed> or <GRCh38_peaks.bed>] -b <homozygous_regions.bed>”. Peaks assigned to hap1 that overlapped homozygous regions were designated as homozygous peaks. Peaks assigned to either haplotype that did not overlap homozygous regions were assigned to hap1 or hap2.

Peaks were lifted over to GRCh38 using custom python scripts. Due to V(D)J recombination in hap2 of NA19240 (**Figure S6** and [36]), peaks were filtered to only include those from position chr14:106037938 (IGHV2-5; GRCh38) to the telomeric end. To assign peaks resulting from the IGH-personalized reference analysis as overlapping low-mappability (mappability < 1) regions, bedtools ‘intersect’ was used with parameters “-wa -f 0.25 -a <heterogygous_peaks.bed> -b <mappability_less_than_1.bed>”.

To determine overlapping intervals, concatenated peaks were input to bedtools ‘merge’ with the options ‘-c 5 -o collapse’ and the output was parsed in R (v4.2.1); upset plots were generated using the ‘ComplexUpset’ package (https://github.com/krassowski/complex-upset) [72].

## Supplemental Figure Legends

**Figure S1. SV regions from which paired-end reads were simulated**

IGV screenshots showing alignment of the IGH locus from the GRCh38 assembly (chr14:105860500-107043718) to hap1 or to hap2 IGH assemblies from NA19240. SV regions from which paired-end reads were simulated are shown.

**Figure S2. Alignment of reads derived from SD sequence (IGHV3-23D) to the NA19240 IGH-personalized reference.**

3,800 paired-end reads (100 bp) simulated from the IGHV3-23D SV (SD) region in hap1 of NA19240 (indicated by blue bar in the “SV regions” track) were aligned to the NA19240 IGH-personalized reference. Simulated reads and hap1 assemblies are both salmon colored. Read mapping results are shown for alignments to the hap1 **(A)** and hap 2 assemblies **(B)**, which number of reads mapped to the IGHV3-23 and IGHV3-23D regions of hap1 and IGHV3-23 region of hap2 indicated.

**Figure S3. Lengths of indicated SV regions in NA19240 IGH haplotypes.**

**Figure S4. Regions of homozygosity in NA12878 and NA19240 IGH haplotypes.**

Lengths of regions of homozygosity in IGH haplotypes (for k-mer=300; see Methods). NA12878 has four regions of homozygosity greater than 15 Kbp in length.

**Figure S5. Peaks assigned to SV regions in IGH-personalized reference peak sets.**

IGV screenshots showing alignment of the IGH locus from the GRCh38 assembly (chr14:105860500-107043718) to hap1 or to hap2 IGH assemblies from NA19240; peaks not lifted over to GRCh38 are within SV regions that are absent from the GRCh38 assembly. (**A**) SV regions on hap1, the hap2 IGH assembly aligned to hap1, homozygous regions, homozygous peaks, and hap1 peaks that were not lifted over to GRCh38. (**B**) SV regions on hap2, the hap1 assembly aligned to hap2, as well as hap2 peaks that were not lifted over to GRCh38.

**Figure S6. NA19240 IGH haplotypes aligned to the GRCh38 reference assembly.**

IGV screenshot showing alignment of NA19240 IGH hap1 and hap2 assemblies to the GRCh38 IGH locus (chr14:105860500-107043718). Sequence lost from hap2 due to V(D)J recombination, as described previously [36], is indicated.

