## Supplementary figures and images for "Addressing technical pitfalls in pursuit of molecular factors that mediate immunoglobulin gene regulation"

### Supplemental Figures

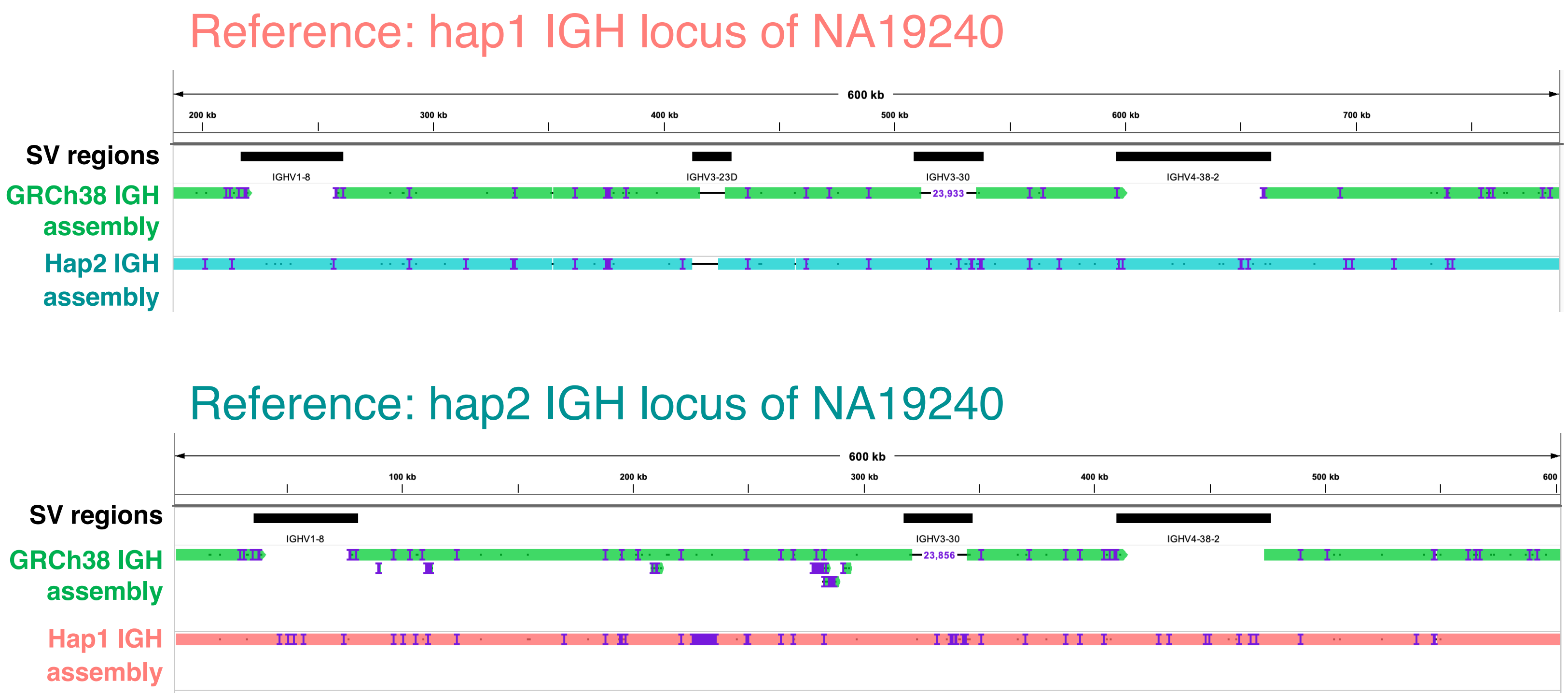

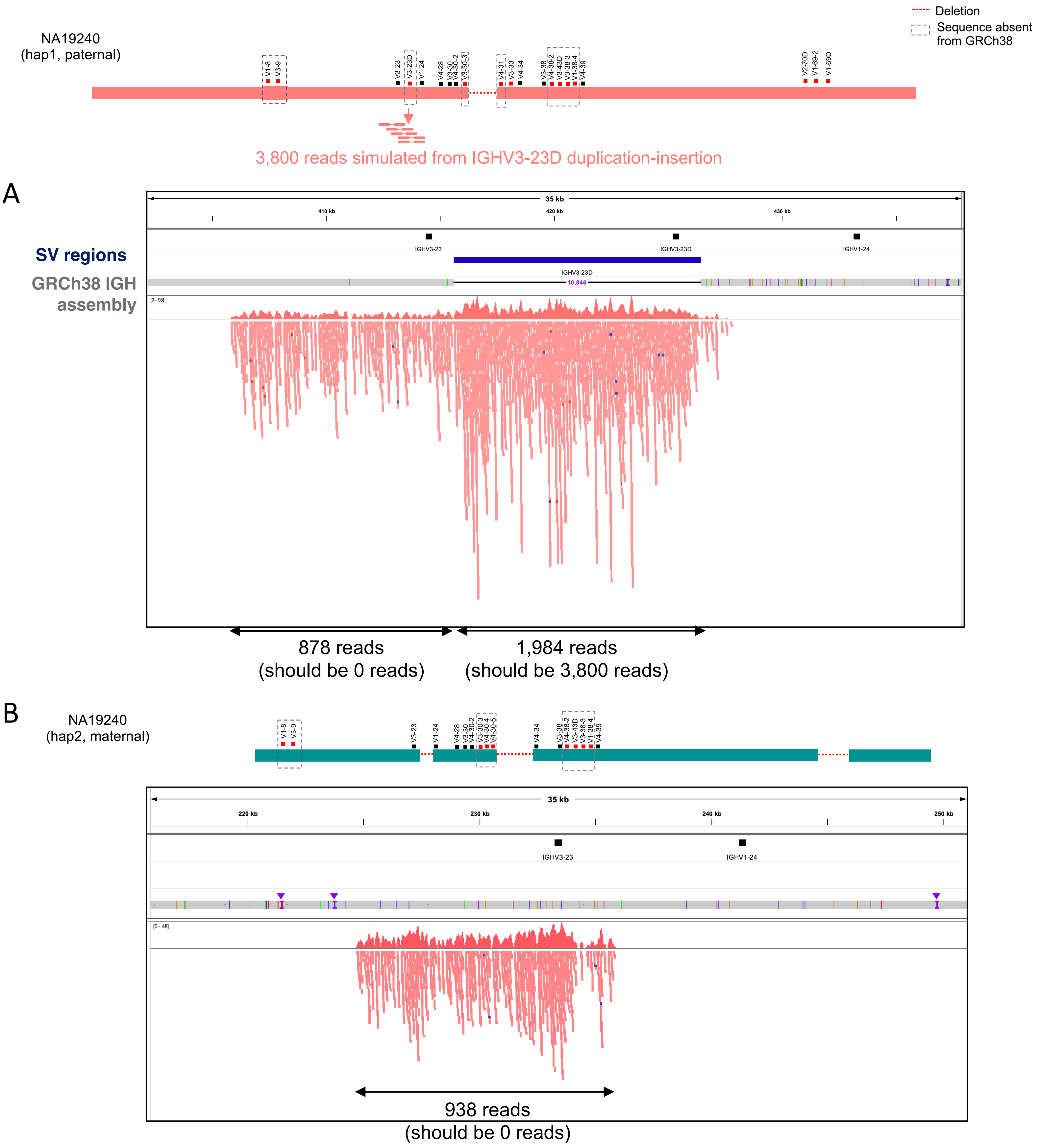

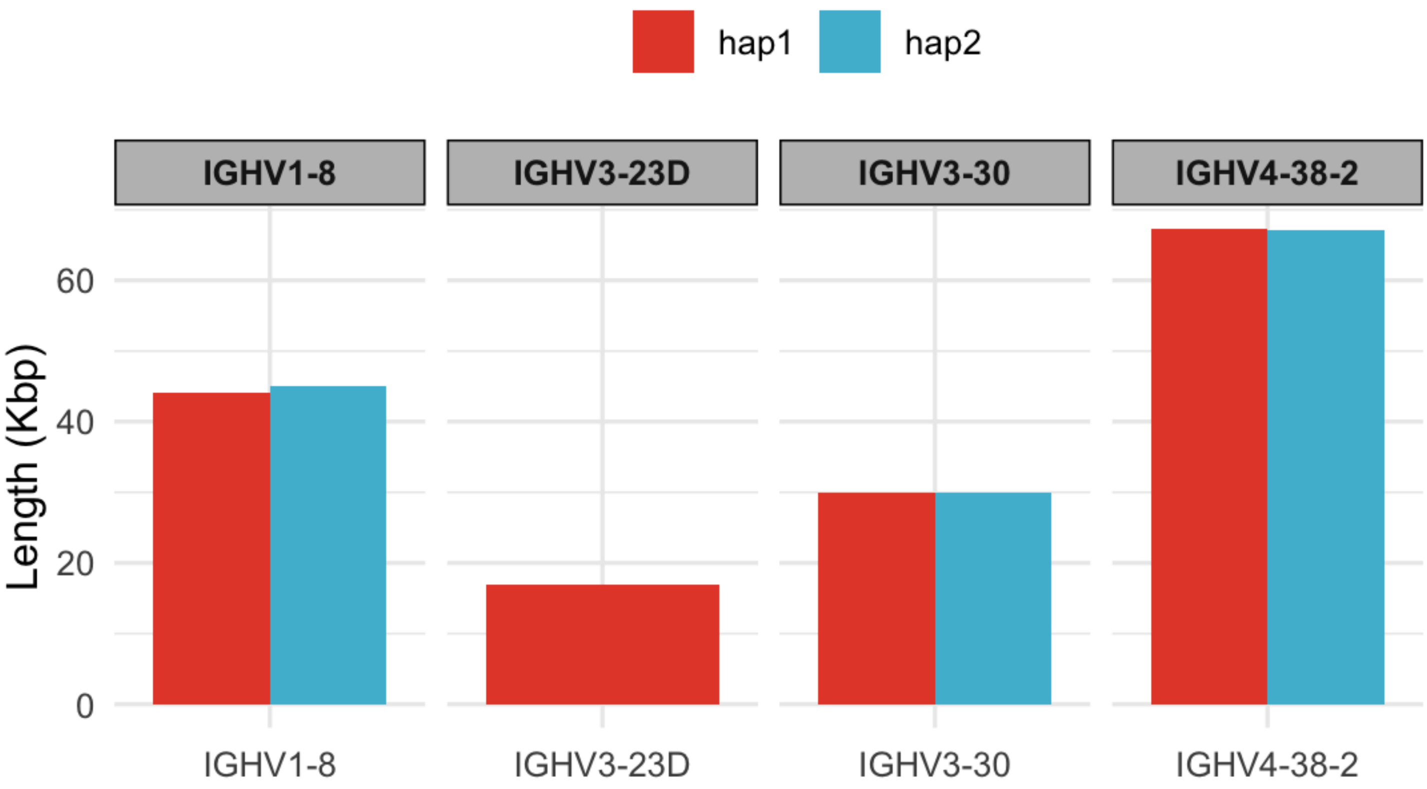

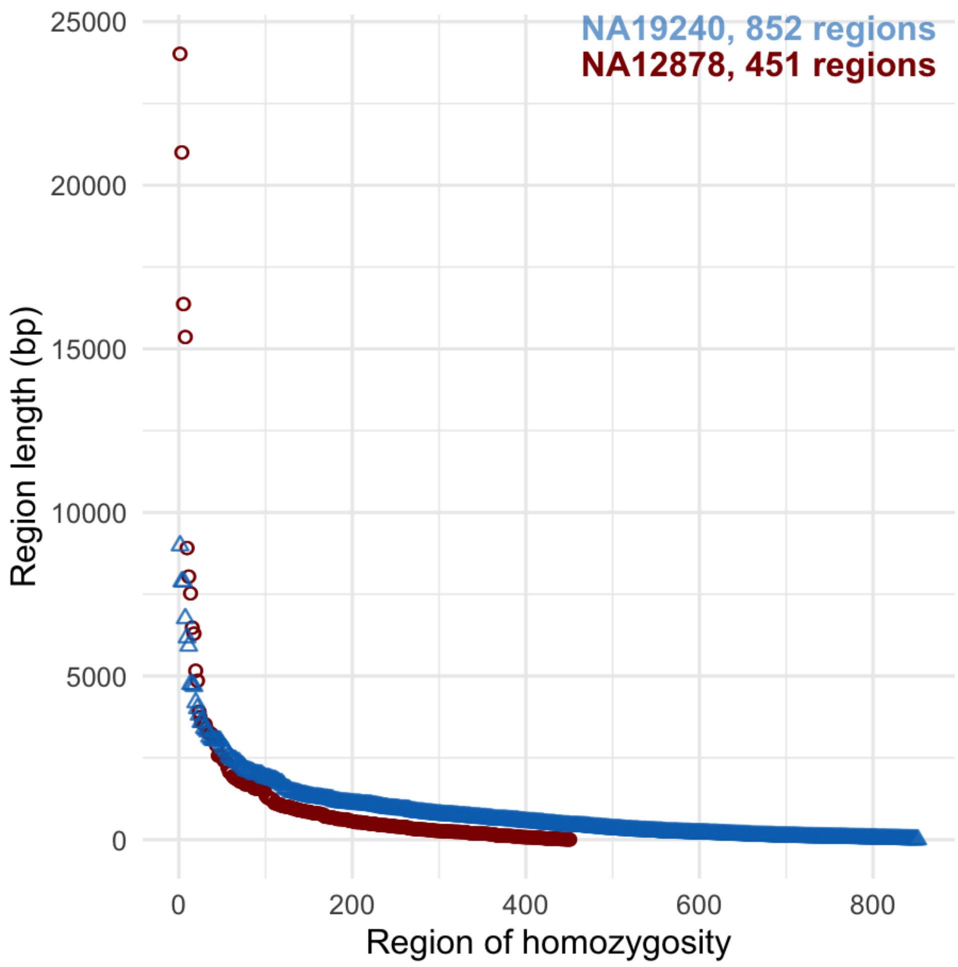

#### Reference: hap1 IGH locus of NA19240

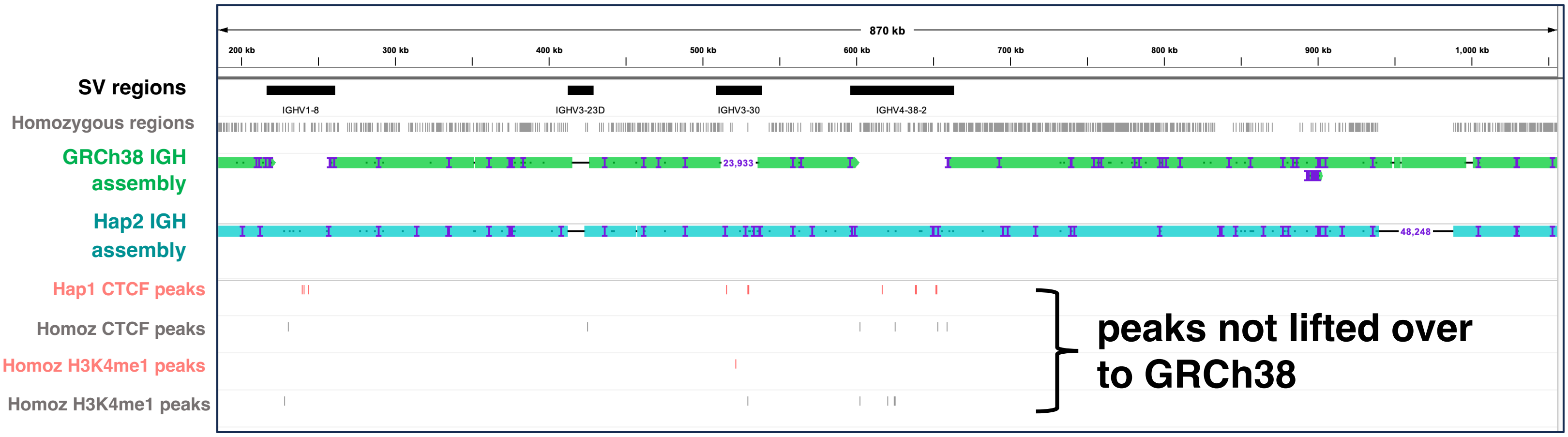


#### Reference: hap2 IGH locus of NA19240

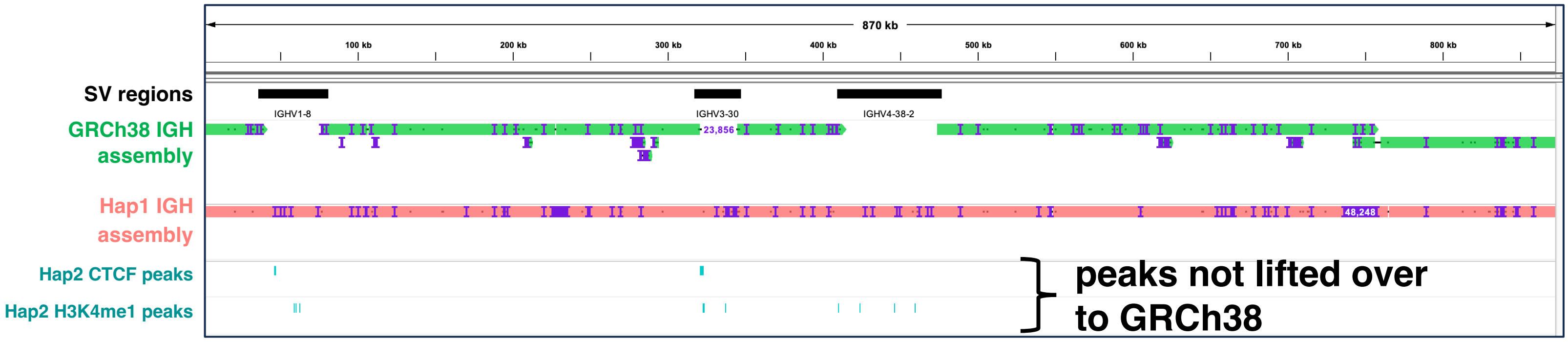


Figure S5

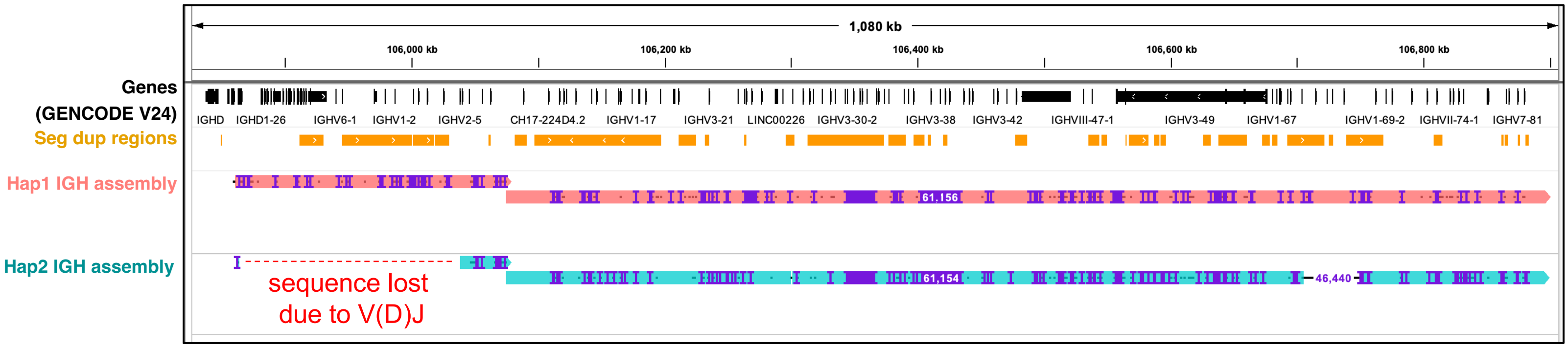
